# New *App* knock-in mice that accumulate wild-type human Aβ as rapidly as *App^NL-G-F^* mice exhibit intensive cored plaque pathology and neuroinflammation

**DOI:** 10.1101/2021.04.30.442122

**Authors:** Kaori Sato, Naoto Watamura, Ryo Fujioka, Naomi Mihira, Misaki Sekiguchi, Kenichi Nagata, Toshio Ohshima, Takashi Saito, Takaomi C. Saido, Hiroki Sasaguri

## Abstract

We previously developed single *App* knock-in mouse models of Alzheimer’s disease (AD), harboring the Swedish and Beyreuther/Iberian mutations with or without the Arctic mutation (*App^NL-G-F^* and *App^NL-F^* mice). These models showed amyloid β peptide (Aβ) pathology, neuroinflammation and cognitive impairment in an age-dependent manner. The former line exhibits extensive pathology as early as 6 months but is unsuitable for investigating Aβ metabolism and clearance because the Arctic mutation renders Aβ resistant to proteolytic degradation and prone to aggregation. In particular, it is inapplicable to preclinical immunotherapy studies due to its discrete affinity for anti-Aβ antibodies. The weakness of the latter model is that it may take as long as 18 months for the pathology to become prominent. We have thus generated a new model that exhibits early deposition of wild-type human Aβ by crossbreeding the *App^NL-F^* line with the *Psen1^P117L/WT^* line. We show that the effects of the pathogenic mutations in the *App* and *Psen1* genes are additive or synergistic. This new mouse model showed more cored plaque pathology and neuroinflammation than *App^NL-G-F^* mice and will help accelerate the development of disease-modifying therapies to treat preclinical AD.

## Introduction

The major pathological hallmark of Alzheimer’s disease (AD), the most common type of dementia, is deposition of amyloid β peptide (Aβ) in the brain^1,2^. Over 300 mutations in the *presenilin 1 (PSEN1)* and *presenilin 2 (PSEN2)* genes and more than 50 mutations in the *amyloid precursor protein (APP)* gene have been identified as disease-associated mutations (Alzforum, http://www.alzforum.org). These findings have led to the development of transgenic mice overexpressing mutant APP or APP/PSEN1 cDNAs^3^. Such mouse models, however, often suffer from experimental limitations caused by overproduction of APP fragments such as C-terminal fragment of APP generated by β-secretase (β-CTF) and APP intracellular domain (AICD), both of which do not appear to accumulate in AD brains and may induce artificial endosomal abnormalities^4^ and transcriptional malfunctions^5^, respectively. Other overexpression artifacts include calpain activation^6^, calpastatin deficiency-induced early lethality^7^ and endoplasmic reticulum stresses^8^. In addition, Gamache *et al*. demonstrated that the random insertion of transgene(s) destroyed unexpectedly large regions in endogenous gene loci of the host animal ^9^. We suggest that all transgenic models overexpressing APP or APP/PSEN1 that are being used in research should be subjected to whole genome sequencing to identify the destroyed loci that may have affected their phenotypes.

To overcome these drawbacks, we previously generated *App^NL-G-F/N-LG-F^* knock-in *(App^NL-G-F^)* and *App^NL-F/NL-F^* knock-in *(App^NL-F^)* mice that harbor the Swedish (KM670/671NL) and Beyreuther/Iberian (I716F) mutations with or without the Arctic (E693G) mutation^3,10^. These mice showed typical Aβ pathology, neuroinflammation and memory impairment^10 11^ and are being used by more than 500 research groups world-wide. Thus far, the *App^NL-G-F^* line has been more frequently used than the *App^NL-F^* line because the former develops Aβ pathology approximately 3 times faster than the latter^10^ and can be conveniently used to analyze downstream events such as neuroinflammation ^12 13 14^, pericyte signalling^15^, oxidative stress ^16 17 18 17^, tau propagation ^19^ and spatial memory impairment ^11 20 21^.

However, the *App^NL-G-F^* line is unsuitable for investigating Aβ metabolism and clearance because the Arctic mutation renders Aβ resistant to proteolytic degradation ^22^ and prone to aggregation ^23^. In particular, it is unsuitable for use in preclinical studies of immunotherapy due to its discrete affinity for anti-Aβ antibodies even in the presence of Guanidine Hydrochloride (GuHCl) ^10^. The Arctic mutation may also interfere with the direct or indirect interactions between Aβ deposition and apolipoprotein E genotype^24^ although there is no experimental evidence. In contrast, the *App^NL-F^* line does accumulate wild-type human Aβ, but it takes as long as approximately 18 months for the pathology to become prominent ^10^: 18 months are too long for students and postdocs to wait.The aim of the present study was thus to generate a new mouse model that accumulates wild-type human Aβ as quickly as the *App^NL-G-F^* model, but without depending on the Arctic mutation.

We devised the strategy to utilize the heterozygous *Psen1^P117L/WT^* mutant line (*Psen1^P117L^*) that exhibited the largest increase in Aβ_42_/Aβ_40_ ratio in the brain among several *Psen1* mutants that we generated ^25^. In the present study, we attempted to crossbreed *App^NL-F^* mice with *Psen1^P117L^* mice despite it being unclear whether their pathogenic effects, both of which act on the γ-cleavage of β-CTF, are additive or not *in vivo*. We demonstrate here that the *Psen1^P117L^* mutation markedly enhances the pathological phenotypes of *App^NL-F^* mice additively or synergistically. We anticipate that these double mutant mice will become highly relevant tools for examining the mechanisms upstream of Aβ deposition and for preclinical screening of disease-modifying therapy candidates without any concern regarding the artificial effect of the Arctic mutation.

## Results

### *App^NL-F^Psen1^P117L^* double-mutant mice produce higher levels of Aβ_42_ than *App^NL-F^* mice

To analyze the combined effect of *App* and *Psen1* mutations on amyloid pathology *in vivo*, we first prepared *App×Psen1* double-mutant mice carrying mutations in the endogenous genes. We crossbred *Psen1^P117L^* mice ^25^, produced by using cytosine base editors ^26^, with *App^NL-F^* mice ^10^ to generate *App^NL-F/NL-F^×Psen1^P117L/WT^* double-mutant mice *(App^NL-F^Psen1^P117L^* mice). It should be noted that the double-mutant mice used in our experiments were heterozygous for the *Psen1* mutation.

*App^NL-F^* and *App^NL-F^ Psen1^P117L^* mice expressed indistinguishable quantities of APP and α/β-CTFs (**Fig. 1a**), suggesting that the P117L mutation does not alter processing of APP by α and β secretases. Consistent with our previous report^10^, the Swedish mutations increased the ratio of β/α-CTFs to an identical extent in both lines. We then quantified Aβ_40_ and Aβ_42_ levels in the cortices of *App^NL-F^* and *App^NL-F^Psen1^P117L^* mice by Enzyme-Linked Immunosorbent Assay (ELISA). At 3 months of age, male *App^NL-F^Psen1^P117L^* mice produced 22.5-fold GuHCl-soluble (Tris-insoluble) Aβ_42_ compared to *App^NL-F^* mice (**Fig. 1b**): female samples showed a similar (26.2-fold) increase. The increase of Aβ_40_ was much smaller, resulting in approximately 11-fold elevation in the Aβ_42_/Aβ_40_ ratio of male *App^NL-F^Psen1^P117L^* mice compared to *App^NL-F^* mice (Fig. 1b). Female mice showed a similar tendency. In 12-month-old *App^NL-F^Psen1^P117L^* mice, the quantity of Aβ_42_ increased considerably in both Tris-soluble and GuHCl-soluble fractions (**Fig. 1c,d**). Given that the 3-month-old single *Psen1^P117L^* mice showed only a 2- to 3-fold increase in Aβ_42_ production compared to wild-type controls ^25^, our data indicate that the combination of the *App^NL-F^* and *Psen1^P117L^* mutations acts on the γ-secretase activity in an additive or synergistic manner.

**Figure 1.**
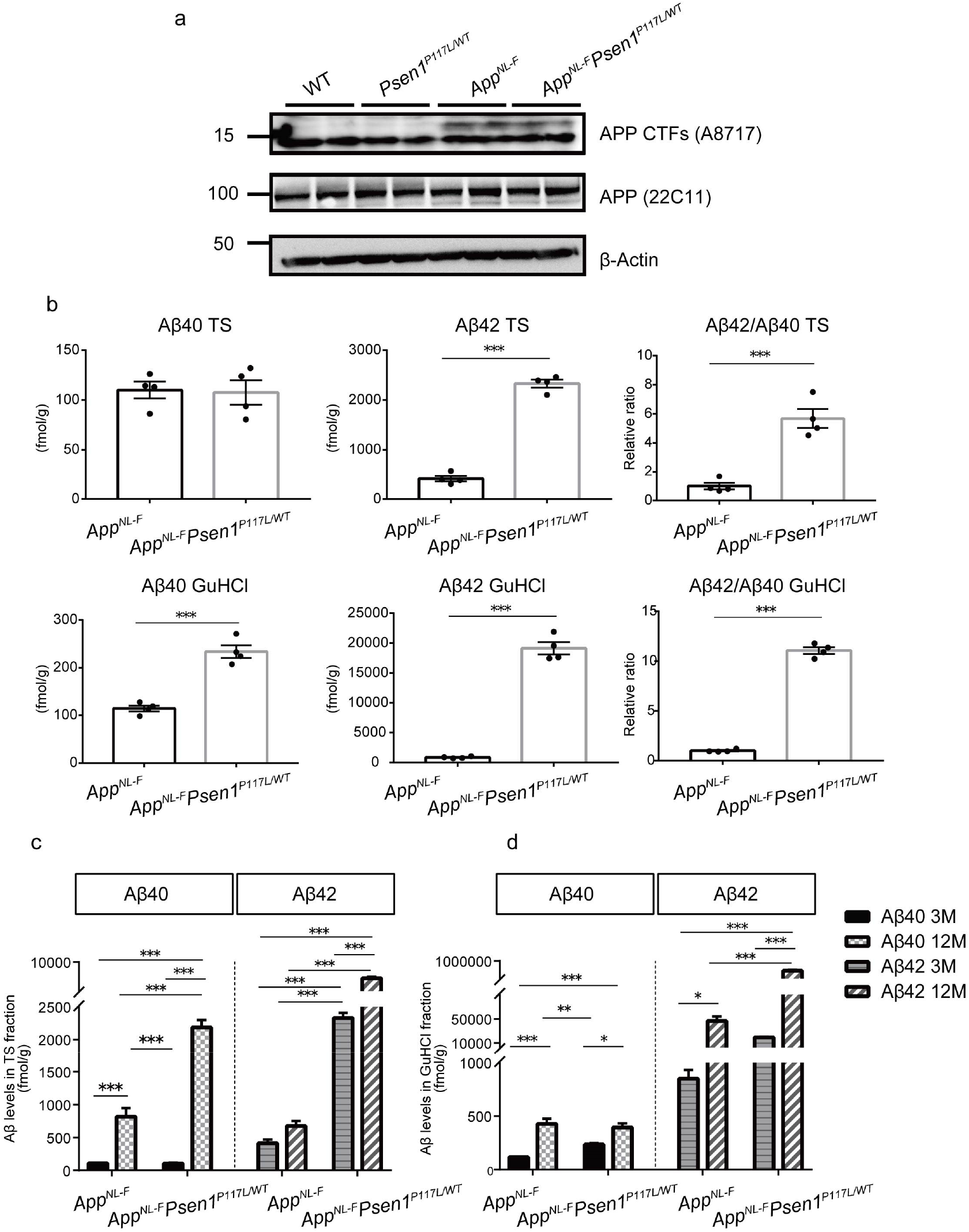
APP processing and Aβ_40_ and Aβ_42_ production in the brains of *App^NL-F^Psen1^P117L^* mice. **(a)**APP processing in the cortices of WT, *Psen1^P117L^, App^NL-F^* and *App^NL-F^Psen1^P117L^* mice. Full blot images of western blotting are shown in Supplementary data 1. (**b)**Aβ_40_ and Aβ_42_ detected by ELISA from the cortices of 3-month-old WT and *App^NL-F^Psen1^P117L^* mice. WT (n=4) and *App^NL-F^Psen1^P117L/WT^* (n=4) (Student’s t-test). (**c, d**) Aβ_40_ and Aβ_42_ using Tris-HCl (**c**) and GuHCl (**d**) soluble fractions from 3- and 12-month-old mice. WT (n=4) and *App^NL-F^Psen1^P117L/WT^* (n=4) (two-way ANOVA followed by Tukey’s multiple comparison test). Each bar represents the mean ± SEM. * P < 0.05, ** P < 0.01, *** P < 0.001.

### The *Psen1^P117L^* mutation also influences Aβ43 production

We previously reported that Aβ_43_ is as pathogenic as Aβ_42_ ^27^. We thus performed Aβ_43_ ELISA on cortices from 3- and 12-month-old *App^NL-F^* and *App^NL-F^Psen1^P117L^* mice. The Tris-soluble, but not insoluble, Aβ_43_ increased more than 2-fold in the brains of *App^NL-F^Psen1^P117L^* mice at 3 months of age compared to *App^NL-F^* mice (**Fig. 2a,b**). Because we treat the “soluble” fractions with GuHCl before the ELISA measurement ^28^, soluble oligomers are likely included in these fractions. Aβ_43_ levels in the GuHCl fractions increased with aging both in *App^NL-F^* and *App^NL-F^Psen1^P117L^* mice (**Fig. 2c**). Some *Psen1* mutations such as I213T and R278I result in the overproduction of Aβ_43_ *in vivo*^29,30^. It is possible that P117L alone or combination with Swedish/Iberian mutations in the *App* gene may lead to an increase in Aβ_43_ by modifying the carboxypeptidase-like activity of γ-secretase in brain ^31,32^. Intriguingly, the Aβ_43_ pathology became more prominent with aging. (See below.)

**Figure 2.**
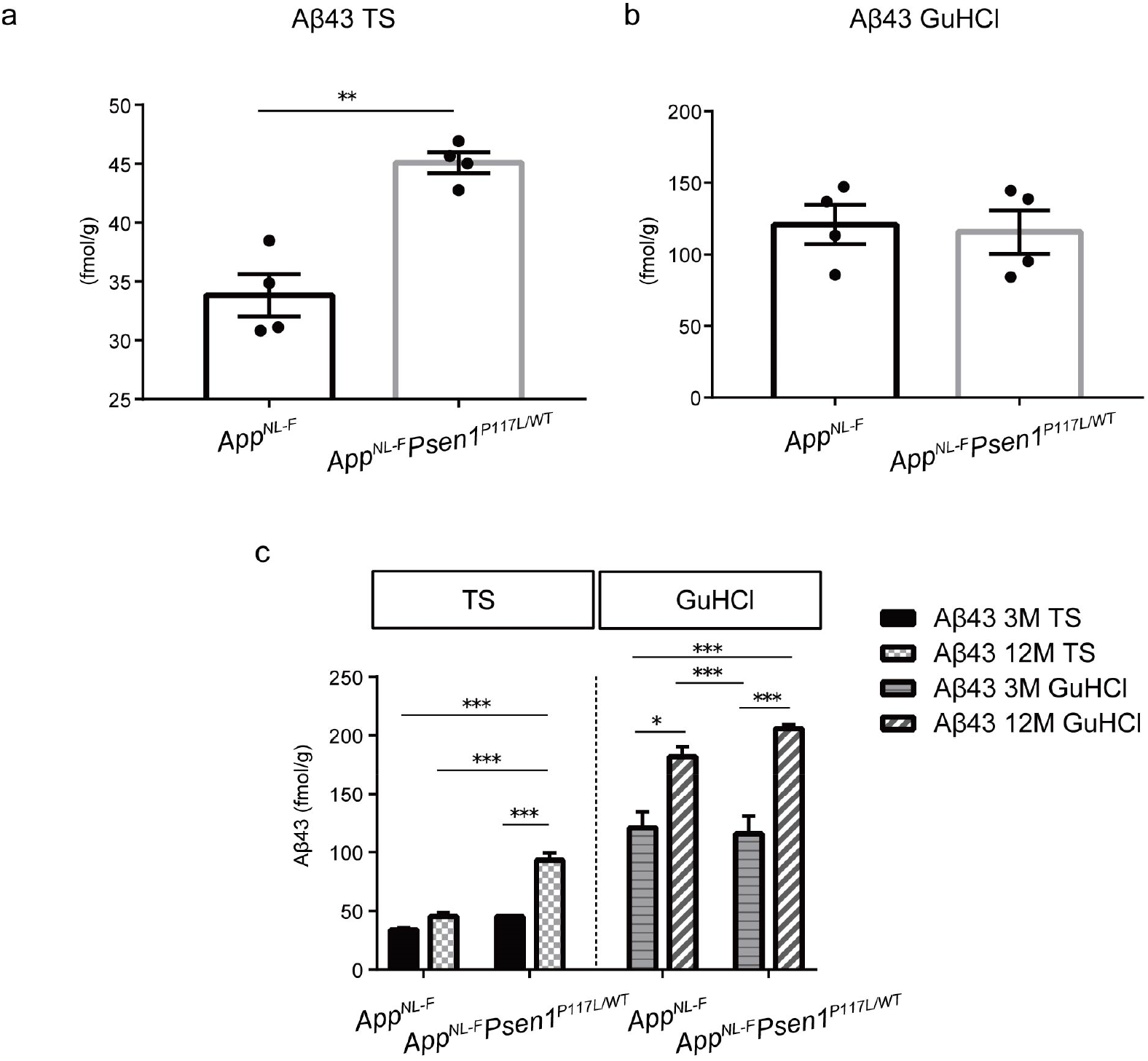
Aβ_43_ levels detected in the cortices of *App^NL-F^* and *App^NL-F^Psen1^P117L^* mice. **(a,b)** Aβ_43_ quantified by ELISA using Tris-HCl (**a**) and GuHCl (**b**) soluble fractions from the cortices of 3-month-old WT and *App^NL-F^Psen1^P117L/WT^* mice. WT (n=4) and *App^NL-F^Psen1^P117L/WT^* (n=4) (Student’s t-test). (**c)**Quantity of Aβ_43_ from Tris-HCl and GuHCl soluble fractions from 3- and 12-month-old mice. WT (n=4) and *App^NL-F^Psen1^P117L/WT^* (n=4) (two-way ANOVA followed by Tukey’s multiple comparison test). Each bar represents mean ± SEM. * P < 0.05, ** P < 0.01, *** P < 0.001.

### Aβ deposition starts as early as 3 months of age in *App^NL-F^Psen1^P117L^* mice

We next examined Aβ pathology in the brains of *App^NL-F^Psen1^P117L^* mice. Immunofluorescence analyses detected Aβ plaques in the cortices of *App^NL-F^Psen1^P117L^* mice at 3 months of age (Fig. 3a and b), whereas *App^NL-F^* mice took as long as 6 months to reach an initial and minimal deposition of Aβ^10^. At 12 months, *App^NL-F^Psen1^P117L^* mice displayed prominent amyloidosis in the cortex and hippocampus comparable to that of *App^NL-G-F^* mice, while significantly fewer Aβ plaques were observed in *App^NL-F^* mice (**Fig. 3a-c**). Of note, the number of subcortical plaques in *App^NL-F^Psen1^P117L^* mice was significantly less than that in *App^NL-G-F^* mice, implying that *App^NL-F^Psen1^P117L^* mice may recapitulate the human pathology in a more faithful manner ^33^. *App^NL-F^Psen1^P117L^* mice produced dominant deposition of Aβ_42_ with minimal Aβ_40_ (**Fig. 3d**), which is consistent with observations made on *App^NL-F^* and *App^NL-G-F^* mice and human samples ^10^. Remarkably, we detected a significantly larger number of Aβ_43_-positive plaques in the cortical, hippocampus and subcortical regions of *App^NL-F^Psen1^P117L^* mice than in those regions of *App^NL-F^* and *App^NL-G-F^* mice (**Fig. 3d,e**). These results imply that the Swedish/Iberian and P117L mutations together may accelerate the generation of longer Aβ species including Aβ_42_ and Aβ_43_, resulting in Aβ pathology at younger ages.

**Figure 3.**
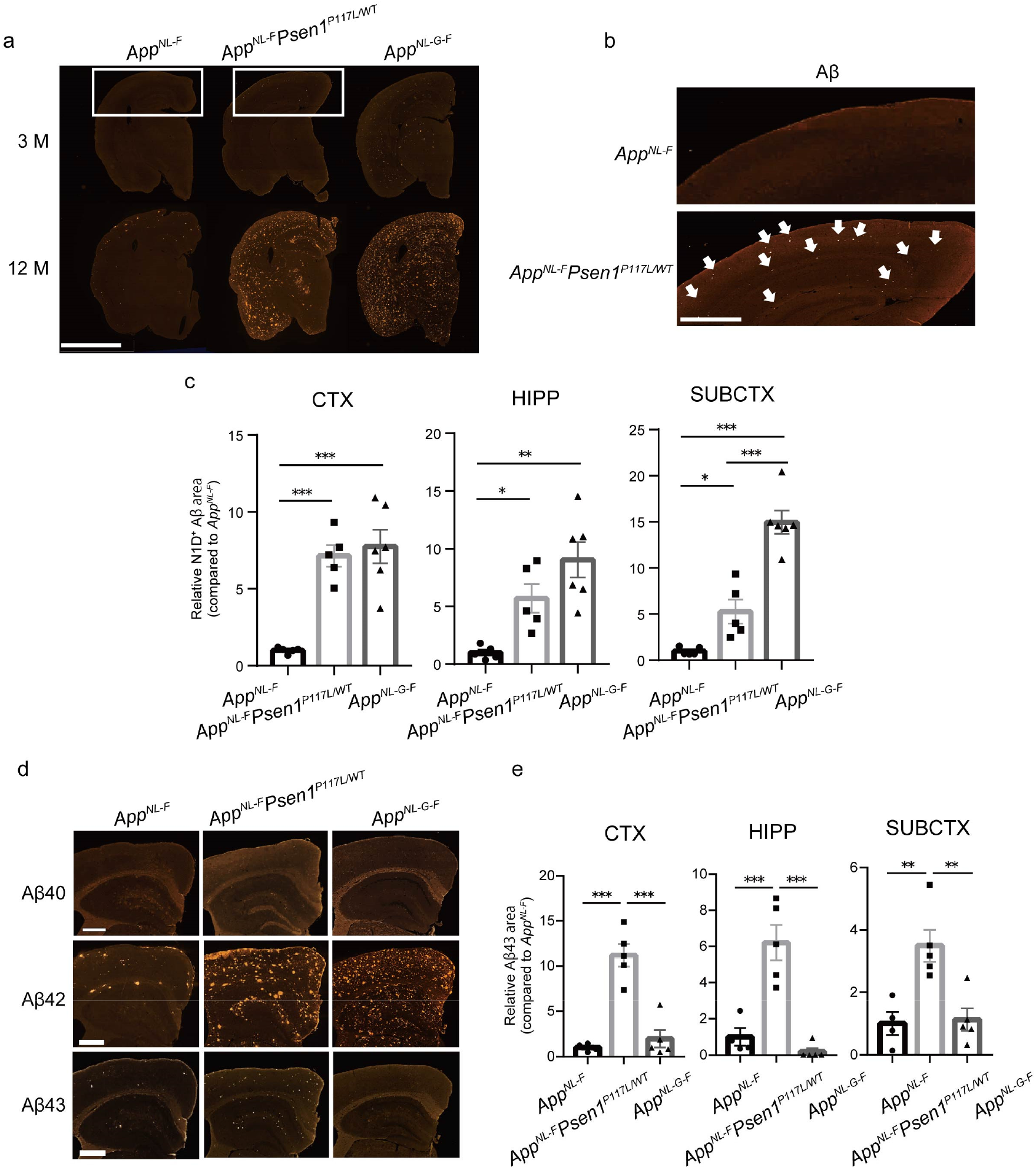
Aβ plaques deposited in the brains of *App^NL-F^Psen1^P117L^* mice. **(a)**Immunofluorescence images showing amyloid pathology in the brains of 3- (top) and 12-month-old (bottom) *App^NL-F^, App^NL-F^Psen1^P117L/WT^* and *App^NL-G-F^* mice. Scale bar represents 2.5 μm. (**b**) High magnification images of tissue sections in (**a**) from 3-month-old animals. Aβ deposition is indicated by white arrows. Scale bar represents 500 μm. (**c)** N1D-positive areas were quantified in cortical (CTX), hippocampal (HIPP) and subcortical (SUBCTX) regions, respectively. *App^NL-F^* (n=5), *App^NL-F^Psen1^P117L/WT^* (n=5) and *App^NL-G-F^* (n=6). (**d)** Sections from 12-month-old mice were immunostained with antibodies specific to Aβ_40_, Aβ_42_ and Aβ_43_. Scale bars represent 50 μm. **(e)**Quantification of Aβ_43_-positive areas in cortical (CTX), hippocampal (HIPP) and subcortical (SUBCTX) regions of 12-month-old mice. *App^NL-F^* (n=4), *App^NL-F^Psen1^P117L/WT^*(n=5) and *App^NL-G-F^* (n=5) (one-way ANOVA followed by Tukey’s multiple comparison test). Each bar represents the mean ± SEM. * P < 0.05, ** P < 0.01, *** P < 0.001.

### Combination of the Swedish/Iberian and P117L mutations is associated with cored Aβ plaque formation

Several lines of evidence support the notion that diversity in Aβ species correlates with plaque morphology such as typical cored plaques ^34–36^. We therefore performed co-staining with N1D antibody raised against Aβ_1-5_ peptide^37^ and 1-Fluoro-2,5-bis(3-carboxy-4-hydroxystyryl)benzene (FSB), which recognizes the β-sheet structure within amyloid fibrils and displays higher fluorescence intensity than 1-bromo-2, 5-bis-(3-hydroxycarbonyl-4-hydroxystyryl)benzene (BSB) and Congo red ^38 39^. We observed that FSB-positive signals were positioned at the center of plaques (**Fig. 4a**) and that the N1D/FSB double-positive plaques were significantly increased in the cortex and hippocampus of *App^NL-F^Psen1^P117L^* mice compared to those of *App^NL-F^* and *App^NL-G-F^* mice (**Fig. 4b**). No significant difference was observed in the subcortical region between *App^NL-F^Psen1^P117L^* and *App^NL-G-F^* mice. The frequent presence of classic dense-cored plaques in the cortex of double-mutant mice was confirmed by 3,3’-diaminobenzidine (DAB) staining (**Fig. 4c**).

**Figure 4.**
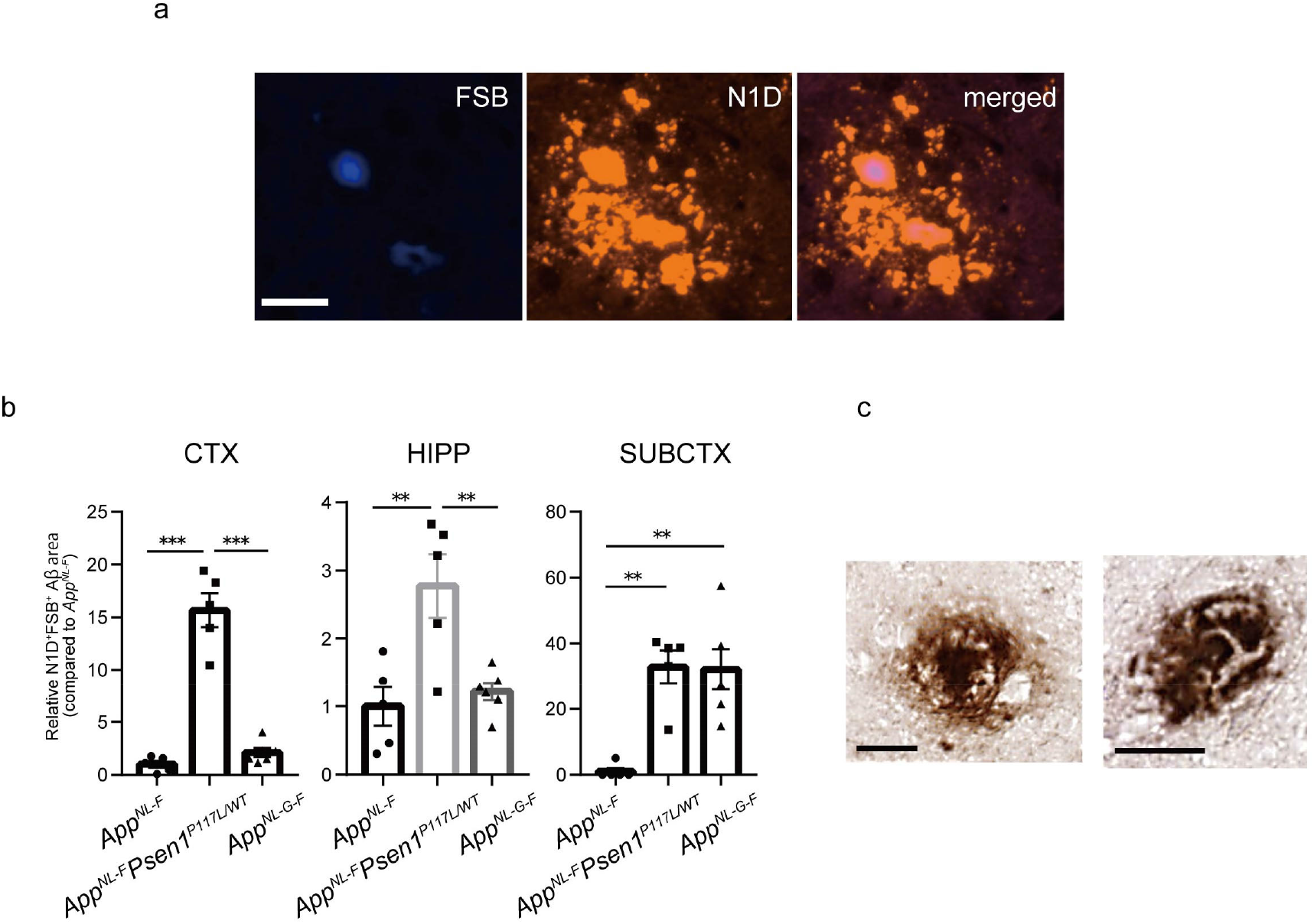
Aβ plaques with a cored structure in *App^NL-F^Psen1^P117L^* mice. **(a)**Brain sections from 12-month-old *App^NL-F^Psen1^P117L/WT^* mice were co-stained with FSB and N1D antibody. (**b)**FSB/N1D double-positive areas were quantified in the brains of *App^NL-F^, App^NL-F^Psen1^P117L/WT^* and *App^NL-G-F^* mice. *App^NL-F^* (n=5), *App^NL-F^Psen1^P117L/WT^* (n=5) and *App^NL-G-F^* (n=6) (one-way ANOVA followed by Tukey’s multiple comparison test). Each bar represents mean ± SEM. * P < 0.05, ** P < 0.01, *** P < 0.001. (**c)**Representative images of dense-core plaques surrounded by a halo effect detected by DAB staining. Scale bars represent 25 μm.

### Neuroinflammation is elevated in *App^NL-F^Psen1^P117L^* mice, particularly in the hippocampus

Neuroinflammation surrounding Aβ plaques manifests as one of the pathological features in AD patients ^10,40,41^, and Genome-Wide Association Studies (GWAS) have suggested etiological involvement of neuroinflammation in AD development^42–44^. We thus analyzed the neuroinflammatory status of three mouse lines (*App^NL-F^, App^NL-F^Psen1^P117L^* and *App^NL-G-F^*) by immunofluorescence using antibodies against Aβ (82E1), microglia (Iba1) and astrocytes (anti-GFAP). We confirmed the presence of glial cells surrounding Aβ plaques in *App^NL-F^Psen1^P117L^* mice (**Fig. 5a,b**). Consistent with our previous reports, single *App^NL-F^* rather than *App^NL-F^* mice exhibit robust microgliosis and astrocytosis accompanying progressive amyloidosis^10,11^. Quantification of immunofluorescence images indicated that more neuroinflammation was evident in *App^NL-F^Psen1^P117L^* and *App^NL-G-F^* than *App^NL-F^* mice (**Fig. 5c**). This was somewhat predictable because *App^NL-F^Psen1^P117L^* and *App^NL-G-F^* mice accumulate more pathological Aβ than *App^NL-F^* mice (**Fig. 3a-c**). A unique observation is that *App^NL-F^Psen1^P117L^* mice exhibited significantly greater microgliosis and astrocytosis than *App^NL-G-F^* mice in the hippocampus despite the indistinguishable levels of Aβ amyloidosis therein. This phenomenon may be associated with increased Aβ_43_ deposition (**Fig. 3d,e**) and cored plaques in *App^NL-F^Psen1^P117L^* mice (**Fig. 4a,b**), but these speculations alone cannot fully account for the neuroinflammation that took place selectively in the hippocampus. In any case, our findings indicate that *App^NL-F^Psen1^P117L^* mice may be suitable for investigating hippocampal neuroinflammation.

**Figure 5.**
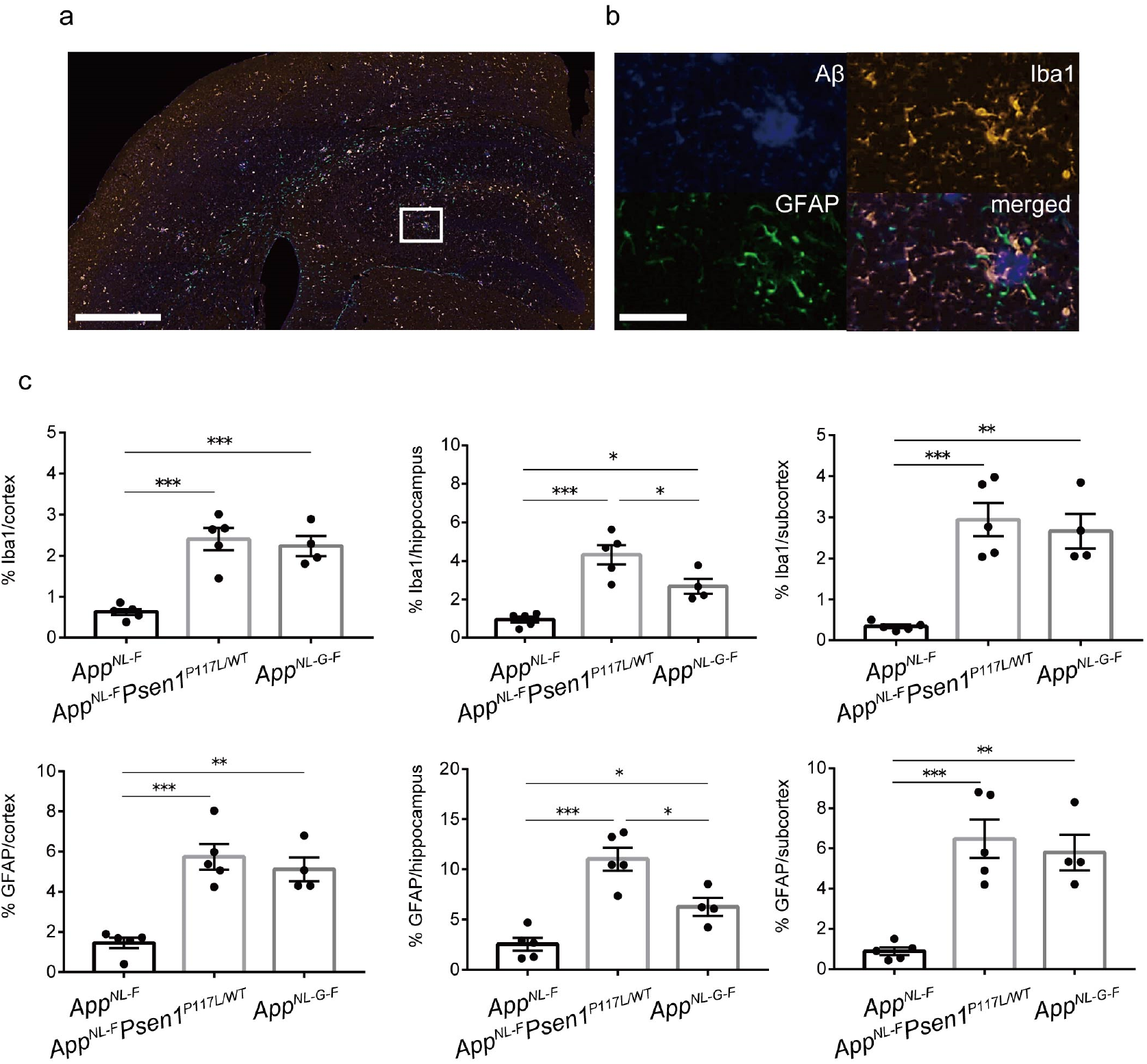
Glial responses in *App^NL-F^Psen1^P117L/WT^* mouse brain. **(a)** Inflammatory signals were detected with GFAP and Iba1 antibodies using brain sections from 12-month-old *App^NL-F^Psen1^P117L/WT^* mice. Aβ pathology was detected by immunostaining with 82E1 antibody. Scale bar represents 500 μm. (**b)** Higher magnification image of the area marked in white in (**a**). Scale bar represents 50 μm. (**c)**Immunoreactive areas of GFAP or Iba1 were quantified in the brains of *App^NL-F^, App^NL-F^Psen1^P117L/WT^* and *App^NL-G-F^* mice. Each bar represents the mean ± SEM. * P < 0.05, ** P < 0.01, *** P < 0.001, *App^NL-F^* (n=5), *App^NL-F^Psen1^P117L/WT^* (n=5) and *App^NL-G-F^* (n=4) (one-way ANOVA followed by Tukey’s multiple comparison test).

## Discussion

The primary aim of the present study was to generate *App* knock-in mice that pathologically accumulate wild-type human Aβ devoid of the Arctic mutation. *App^NL-G-F^* mice carrying the Arctic mutation exhibit the most rapid and aggressive pathology among all the *App* knock-in mice that had been created. Our motivation was based on the undesirable nature of the Arctic mutation that impedes the physiological metabolism and clearance of Aβ^22,23^, thus making it difficult to study etiological processes upstream of Aβ deposition. Arctic Aβ also binds to anti-Aβ antibodies raised against Aβ peptide in a distinct manner ^10^, making *App^NL-G-F^* mice unsuitable for preclinical immunotherapy studies. The generation of *App* knock-in mice that accumulate wild-type human Aβ without the Arctic mutation as quickly as *App^NL-G-F^* mice is therefore a prerequisite before disease-modifying strategies targeting mechanisms upstream of Aβ deposition can be developed.

Introduction of the *Psen1^P117L^* mutation into *App^NL-F^* mice resulted in an unexpected acceleration of Aβ_42_ and Aβ_43_ deposition. Although the numbers of cortical and hippocampal plaques visualized by an antibody specific to the N-terminus of Aβ (N1D) were indistinguishable between *App^NL-G-F^* and *App^NL-F^Psen1^P117L^* mice, *App^NL-F^Psen1^P117L^* mice showed a larger number of cored plaques in the cortex and hippocampus, and more gliosis in the hippocampus than was seen in *App^NL-G-F^* mice. Mechanisms accounting for these observations are unclear and beyond the scope of the present study, but *App^NL-F^Psen1^P117L^* mice may become a useful tool for examining the roles of hippocampal neuroinflammation in the etiology of AD. Other groups have also attempted to combine the *App* and *Psen1* mutations. Flood *et al.* generated double knock-in mice that harbored the Swedish mutations in the *App* gene and the P264L/P264L mutation in the *Psen1* gene ^45^. These mice showed elevation of Aβ_42_ levels and pathological amyloidosis without overexpression of APP. Although this model has seldom been used in the research community presumably because the progression of pathology was too slow and mild, both their group and ours share similar ideas and goals. Similarly, Li *et al.* generated a mouse model of cerebral amyloid angiopathy (CAA)^46^. Unlike the models generated by Flood *et al.* and Li *et al.*, our model mice are heterozygous for the *Psen1* mutation, making them easier to breed.

We must however point out that *App^NL-F^Psen1^P117L^* mice are probably inadequate for studying β- and γ-secretases and their modifiers because the cleavages catalyzed by these secretases are artificially altered by the mutations. We thus expect the mutant mice to become more suitable for the examining catabolism and clearance of Aβ than its anabolism. We will share these mice with the AD research community to accelerate the fight against a disease that deprives patients of their human dignity.

## Author contributions

KS, NW, KN, TS, TCS and HS designed the research plan. KS, NW, RF, NM and HS performed the experiments. KS, NW, KN, TS, TCS and HS analyzed and interpreted data. KS, NW, TCS and HS wrote the manuscript together. TO, TCS and HS supervised the entire research.

## Acknowledgements

We thank Yukiko Nagai-Watanabe for secretarial work. This work was supported by AMED under Grant Number JP20dm0207001 (Brain Mapping by Integrated Neurotechnologies for Disease Studies (Brain/MINDS)) (TCS).

## Materials and Methods

### Animals

*App^NL-F^* and *App^NL-G-F^* mice were described previously ^10^. *Psen1^P117L^* mice ^25^ were crossed with the *App^NL-F^* mice to generate *App/Psen1* double mutant mice. All double mutant mice used in this study were homozygous for the *App* mutations and heterozygous for the *Psen1* mutation (*App^NL-F^Psen1^P117L^*). C57BL/6J mice were used as controls. Male mice were used for biochemical analyses and both male and female mice were used for immunohistochemical studies. All mice were bred and maintained in accordance with regulations for animal experiments promulgated by the RIKEN Center for Brain Science.

### Genotyping

Genomic DNA was extracted from mouse tails in lysis buffer (10 mM pH 8.5 Tris-HCl, 5 mM pH 8.0 EDTA, 0.2% SDS, 200 mM NaCl, 20 μg/ml proteinase K) and subjected to PCR, followed by Sanger sequencing analysis. Primers used for genotyping have been described previously ^10,25^.

### Brain sample preparation

Mice were anesthetized with isoflurane, transcardially perfused and fixed with 4% paraformaldehyde in PBS. The brains were dissected on ice into two halves at the midline. One hemisphere was divided into several parts and stored at −80 °C for biochemical analysis, while the other was incubated at 4 °C for 24 h and rinsed with PBS until paraffin processing for histochemical analysis.

### Western blotting

Mice brain tissues were homogenized in lysis buffer [50 mM Tris pH 7.6, 0.15 M NaCl and Complete protease inhibitor cocktail (Roche)]. Homogenates were incubated at 4 °C for 1 h and centrifuged at 15000 rpm for 30 min, and the supernatants were collected as loading samples. Equal amounts of proteins per lane were subjected to SDS-PAGE and transferred to PVDF or nitrocellulose membranes (Invitrogen). To detect APP-CTFs, delipidated samples were loaded and the membrane was boiled for 5 min in PBS before the next step. After washing and blocking at room temperature, the membranes were incubated at 4 °C overnight with primary antibodies against APP (1:1000, Millipore) or APP-CTFs (1:1000, Sigma-Aldrich), or against β-Actin as a loading control (1:5000, Sigma). The target protein on the membrane was visualized with ECL Select (GE Healthcare) and a Luminescent Image Analyzer LAS-3000 Mini (Fujifilm).

### Immunostaining

Paraffin-embedded mouse brain sections were subjected to deparaffinization and then antigen retrieval was performed by autoclave processing at 121 °C for 5 min. After inactivation of endogenous peroxidases using 0.3%H2O2 solution for 30 min, the sections were washed with TNT buffer (0.1 M Tris pH 7.5, 0.15 M NaCl, 0.05% Tween20), and blocked for 30 min in TNB buffer (0.1 M Tris pH 7.5, 0.15 M NaCl) and incubated in the same buffer with primary antibodies at 4 °C overnight. The primary antibody dilution ratios were as follows: Aβ_40_ (1:100, IBL), Aβ_42_ (1:100, IBL), Aβ_43_ (1:50, IBL), Aβ1-5 (N1D)^37^ (1:200), N-terminus of Aβ(82E1) (1:500, IBL), GFAP (1:200, Millipore) and Iba1 (1:200, Wako). Amyloid pathology was detected using biotinylated secondary antibody and tyramide signal amplification as described previously ^47^. For detection of glial activation, secondary antibodies conjugated with Alexa Fluor 488 or 555 were used. Before mounting, the sections were treated when necessary with DAPI diluted in PBS. Data images were obtained using a NanoZoomer Digital Pathology C9600 (Hamamatsu Photonics). Immunoreactive signals were quantified by Definiens Tissue Studio (Definiens).

### DAB staining

Targeted signals were detected and visualized using VECTASTAIN *Elite* ABC Rabbit IgG kit (Funakoshi) and DAB • TRIS tablets (Mutokagaku). After deparaffinization and antigen retrieval treatment of mouse brain sections, endogenous peroxidases were inactivated using 0.3% H_2_O_2_ solution for 30 min. The sections were blocked with 3 drops of goat serum in PBS for 30 min at room temperature and incubated with N1D antibody at 4 °C overnight. The sections were washed with PBS and incubated with the *Elite* ABC solution for 30 min and subsequently stained with DAB solution following the manufacturer’s instructions. Before mounting, dehydration treatment was performed.

### FSB staining

The PFA-fixed tissue sections were deparaffinized, incubated in 0.01%FSB solution in EtOH for 30 min and then rinsed in saturated Li_2_Co_3_ in water for 15-20 sec at room temperature. The sections were differentiated in EtOH for 3 min followed by immersion in water for 5 min to stop the reaction. Readers should refer to the *Immunostaining* section concerning methods for subsequent treatments following antigen retrieval.

### ELISA

Mouse cortical samples were homogenized in buffer A (50 mM Tris-HCl, pH 7.6, 150 mM NaCl and protease inhibitor cocktail) using a medical beads shocker. The homogenized samples were directed to centrifugation at 70000 rpm for 20 min at 4 °C, and the supernatant was measured and collected as a Tris-soluble (TS) fraction in 6 M guanidine-HCl (Gu-HCl) solution containing 50 mM Tris and protease inhibitors. The pellet was loosened with the buffer A and centrifuged at 70000 rpm for 5 min at 4 °C, and then dissolved in 6 M Gu-HCl buffer. After incubation at room temperature for 1 h, the sample was sonicated at 25 °C for 1 min. Subsequently, the sample was centrifuged at 70000 rpm for 20 min at 25 °C and the supernatant collected as a Gu-HCl fraction. 100 μl of TS and Gu-HCl fractions were loaded onto 96-well plates and incubated at 4 °C overnight using the Aβ_40_, Aβ_42_ and Aβ_43_ ELISA kit (Wako) according to the manufacturer’s instructions.

### Statistics

All data are presented as the mean ± S.E.M. within each figure. For comparisons between two groups, data were analyzed by Student’s *t*-test. For comparisons among more than three groups, we used one-way analysis of variance (ANOVA) followed by Dunnett’s post hoc analysis or Tukey’s post hoc analysis. Statistical analyses were performed using GraphPad Prizm 8 software (GraphPad software). The levels of statistical significance were shown as *P*-values: * *P*< 0.05, ** *P*< 0.01, *** *P*< 0.001.

## Supplementary Figures

**Supplementary Figure 1.**
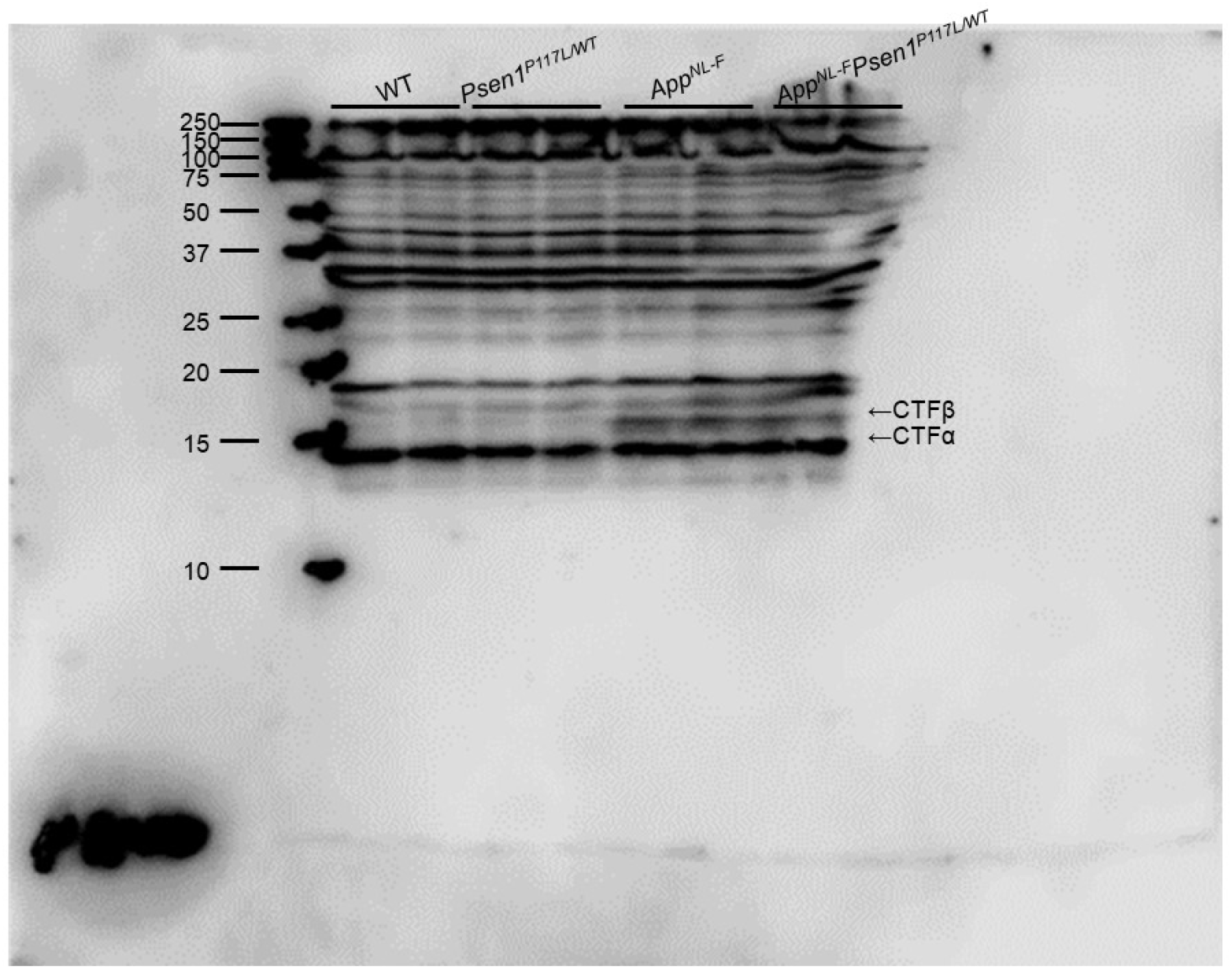
Full Western blots of CTFs shown in Figure 1a.

**Supplementary Figure 2.**
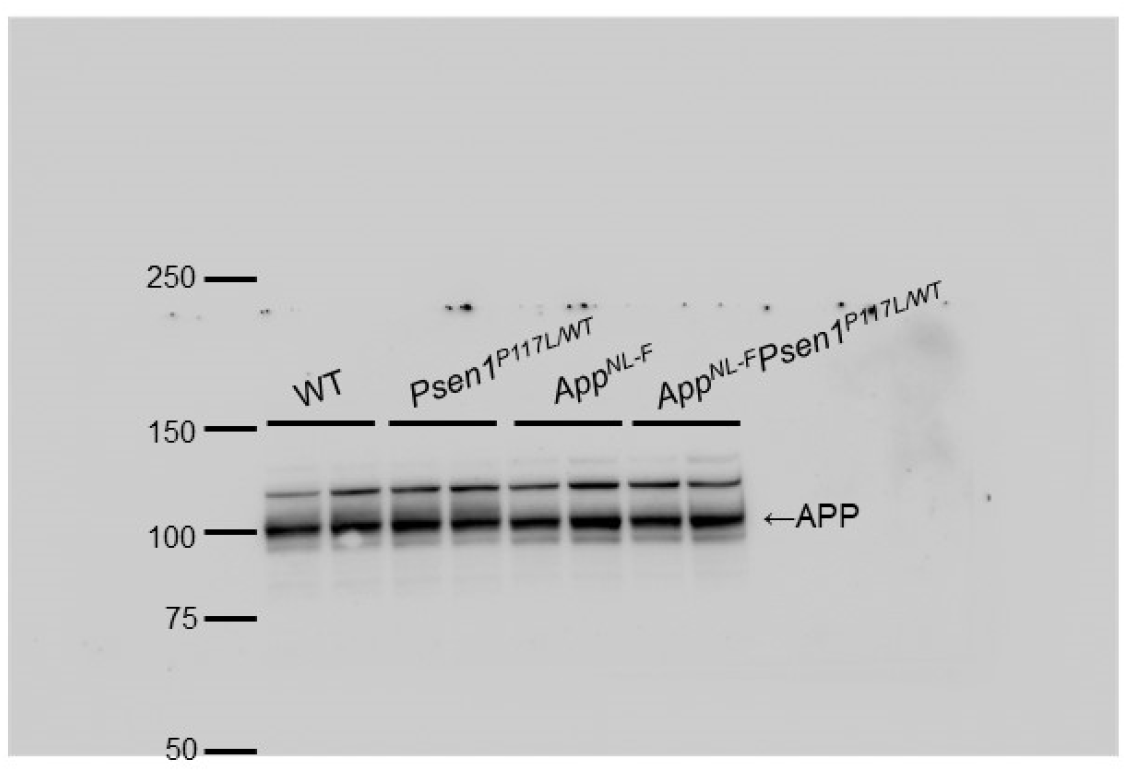
Full Western blots of APP shown in Figure 1a.

